# What kind of maternal effects are selected for in fluctuating environments?

**DOI:** 10.1101/034546

**Authors:** S. R. Proulx, H. Teotónio

## Abstract

Adaptation to temporally fluctuating environments can be achieved through evolution of fixed genetic effects, by phenotypic plasticity (either developmental plasticity or trans-generational plasticity), or by randomizing offspring phenotypes (often called diversifying bet-hedging). Theory has long held that plasticity can evolve when information about the future environment is reliable while bet-hedging can evolve when mixtures of phenotypes have high average fitness (leading to low among generation variance in fitness). To date, no study has studied the evolutionary routes that lead to the evolution of randomized offspring phenotypes on the one hand or deterministic maternal effects on the other. We develop simple, yet general, models of the evolution of maternal effects and are able to directly compare selection for deterministic and randomizing maternal effects and can also incorporate the notion of differential maternal costs of producing offspring with alternative phenotypes. We find that only a small set of parameters allow bet hedging type strategies to outcompete deterministic maternal effects. Not only must there be little or no informative cues available, but also the frequency with which different environments are present must fall within a narrow range. By contrast, when we consider the joint evolution of the maternal strategy and the set of offspring phenotypes we find that deterministic maternal effects can always invade the ancestral state (lacking any form of maternal effect). The long-term ESS may, however, involve some form of offspring randomization, but only if the phenotypes evolve extreme differences in environment-specific fitness. Overall we conclude that deterministic maternal effects are much more likely to evolve than offspring randomization, and offspring randomization will only be maintained if it results in extreme differences in environment-specific fitness.

## 1 Introduction

Variability in the environment is ubiquitous and is expected to provide a significant opportunity for selection (Proulx and Phillips, 2005). When the environment fluctuates on a time scale of generations or longer, then one route for adaptation is the production of discrete phenotypes that vary in their performance over the range of environments that the population typically experiences. A long-standing theoretical dichotomy exists between strategies that produce a fixed set of phenotypes (Crean and Marshall, 2009; Kaplan and Cooper, 1984; Seger and Brockman, 1987; Bull, 1987) and strategies that allow some form of phenotypic plasticity (Hoyle and Ezard, 2012; Kuijper and Hoyle, 2015; Kuijper et al., 2014; Tufto, 2015; Donaldson-Matasci et al., 2013, 2010; Jablonka et al., 1995; Leimar and McNamara, 2015; Donaldson-Matasci et al., 2010). In general, such strategies can be considered a form of phenotypic plasticity in that a single genotype can lead to adults with distinct phenotypes. In scenarios where the environment fluctuates over generations, fitness can be approximately measured as the geometric mean of a genotypes reproductive output (Cohen, 1966; Seger and Brockman, 1987; Proulx and Adler, 2010; Saether and Engen, 2015). While the geometric mean fitness concept depends on the frequency of environment types, and not on the sequence of environmental transitions (Seger and Brockman, 1987), the fitness consequences of a plastic strategy may actually depend on the order of environmental transitions. For example, under an epigenetic “phenotypic memory” model, the fitness of a strategy depends on the probability that a parent has a phenotype that matches the parental environment and that their offspring then experience the same environment (Jablonka et al., 1995). Under strict developmental phenotypic plasticity, the adult (i.e. fitness related) phenotype is determined by the offspring genotype and juvenile environment. If the juvenile environment is a good predictor of the adult environment (i.e. there is high mutual information entropy between juvenile and adult environment) then a developmentally plastic genotype may have high fitness (Simons, 2011).

Seger and Brockman (Seger and Brockman, 1987) attempted to clarify then common misconceptions about reproductive variance, risk, and population genetic change. They argued that the term “bet hedging” should be reserved for situations where a change in genotype increases the geometric mean fitness at the expense of a decrease in the mean of fitness. In this framework, bet hedging can be accomplished either by producing a fixed phenotype that is insensitive to environmental variability (termed conservative bet hedging) or by producing a range of phenotypes at random such that the ensemble of offspring produced has a higher geometric mean fitness (termed diversifying bet hedging). Diversifying bet hedging strategies can achieve high geometric mean fitness by producing a mixture of phenotypes that vary in their environment-specific performance. This type of strategy always involves a certain amount of waste because a fraction of offspring have phenotypes that do not match the environment. This potential fitness can be quantified in an information theoretic sense and used to understand how strong selection can be to take advantage of cues that predict the future environment (Donaldson-Matasci et al., 2013, 2010).

Given the diversity of work on the evolution of developmental plasticity, maternal effects, and phenotypic diversification, our contribution aims to identify the environmental conditions and phenotypic trade-offs that lead to the evolution of particular forms of maternal effect. We start by considering scenarios where the same phenotypic trade-offs apply to genotypes that produce phenotypically diversified offspring as apply to genotypes that influence offspring phenotype through environment-specific maternal effects. While prior work has identified conditions that favor maternal effects on the one hand, and conditions that favor diversifying bet-hedging on the other, our approach allows us to consider both in a common framework and determine which type of maternal effect is most likely to evolve and be maintained.

Our goal here is to understand how maternal strategies can evolve in a fluctuating environment when there is little opportunity for developmental plasticity. It is generally understood that developmental plasticity can evolve when individuals have access to reliable information early in development that can predict their future environment, and when the future environment is relatively constant for the adult lifespan (Uller, 2008). However, in many cases the timing of development and environmental exposure makes it difficult for a developing offspring to independently acquire and utilize this sort of information. This would be the case, for instance, if juvenile survivorship depends on phenotypes that must be present before the developing individual is able to express them. We recently showed this to be the case for survivorship of *C. elegans* when embryos are exposed to anoxia (Dey et al., 2015).

In scenarios where mothers can determine offspring phenotype the range of maternal strategies encompasses mechanisms that produce randomized offspring and those that represent anticipatory maternal effects. Mechanistically, the maternal strategies in question can operate by deterministically altering offspring phenotype in response to maternal environment/phenotype (**Deterministic Maternal Effect**), or by randomizing offspring phenotype (**Randomized Maternal Effect**). We use these terms since they describe specific mechanistic stratagies without implying a specific fitness benefit or selective outcome (as bet hedging and anticipatory maternal effects do). A further extension is diversifying bet hedging around a norm of reaction where a genotype uses some environmental cues to shift the probabilistic distribution from which phenotypes are drawn (Crean and Marshall, 2009; Furness et al., 2015). We refer to situations where environmental cues are used to alter the distribution of offspring phenotypes as a **Hybrid Maternal Effect**.

Here we focus on situations where the environment fluctuates following a Markovian stochastic process and assume that developing offspring do not have direct access to environmental information themselves. We focus on the situation where mothers are able to influence offspring phenotype and take a general approach to the form of the maternal effect. Much prior work on the evolution of maternal effects and phenotypic variance make vague assumptions about how phenotype is mechanistically determined and ignore trade-offs that might underlie the production of diversified broods. We investigate a fairly general scenario whereby mothers are able to directly influence offspring phenotype, both by investing resources and by directly affecting development of offspring (i.e. through maternally transferred resources and RNA). We consider a scenario where the maternal strategy involves production of a pair of offspring phenotypes with a maternally determined probability that depends on the environment that the mother experienced. Given this range of maternal effects, we explore the joint evolution of the maternal strategy and set of possible offspring phenotypes. The range of maternal strategies includes the notion of “diversified bet hedging”, where mothers produce a range of offspring phenotypes, as well as deterministic maternal effects, where mothers produce offspring of a particular phenotype whenever the mother encounters a specific environment. We also investigate intermediate strategies that allow randomization of offspring phenotype in an environment-specific manner.

We take several approaches to understanding this problem, but our overall approach aims to understand the long term population genetic outcomes when recurrent mutation introduces mutants that affect both the maternal effect strategy and the phenotypes that make up that strategy. There are many ways that theorists have approached such long term evolutionary dynamics that include Gillespie’s *Strong Selection Weak Mutation* (SSWM) approach (Gillespie, 1991), Hammerstein’s “streetcar theory” (Hammerstein, 1996), and the Adaptive Dynamics approach (Dieckmann and Law, 1996; Champagnat et al., 2001; Proulx and Day, 2002). We take a hybrid approach and first explore the population genetics of segregating maternal effect mutants, and then study the joint evolution of maternal effect strategies with offspring phenotypes. This allows us to examine the origin of maternal effect strategies from ancestral populations that have already experienced phenotypic evolution with purely genetic inheritance.

Our main conclusion is that deterministic maternal effects can be beneficial in a wide range of conditions, and are able to evolve *de novo* from any ESS that lacks maternal effects. In contrast, bet-hedging is only beneficial when there is already a strong fitness effect of the alternative phenotypes, and experiences no or weak selection from ESS that lacks maternal effects. Nevertheless, bet-hedging may evolve as a derived feature in populations that have already evolved deterministic maternal effects, but only when it is possible to evolve dichotomous phenotypes that have extreme differences in environment-specific fitness.

| Variable | Definition |
| --- | --- |
| $\rho_{i \rightarrow j}$ | the probability that the environment changes from $i$ to $j$ between two generations |
| $\alpha$ | The frequency of generations that experience environment 1 |
| $w_i(x)$ | the offspring fitness of genotype $x$ in environment $i$ |
| $c(x)$ | the fecundity of advantage of a female producing all offspring of phenotype $x$ |
| $\delta_i$ | the relative fitness of phenotype 2 in environment $i$ |
| $\varsigma$ | the relative fecundity of a female who produces only phenotype 2 |
| $\gamma_i$ | the probability that a mother produces offspring with phenotype 1 when she experiences environment $i$ |
| $w_{i,j}(x)$ | the fitness of a mother with genotype $x$ making the transition from environment $i$ to environment $j$ |
| $\hat{w}(x)$ | the geometric mean fitness of genotype $x$ |
| $z_i$ | The $i$th phenotypic trait value produced by a maternal effect strategy |
| $\sigma_i$ | The Gaussian parameter for the strength of selection in environment $i$ |
| DME | Deterministic Maternal Effect, where mothers determine offspring phenotype based on their experience |
| RME | Randomized Maternal Effect, where mothers produce a probabilistic distribution of offspring phenotypes irrespective of their experience |
| HME | Hybrid Maternal Effect, where mothers produce a probabilistic distribution of offspring phenotypes based on their experience |

## 2 The Models

### 2.1 Environmental Fluctuations

We consider a scenario where the environment fluctuates via a stochastic process that may have autocorrelation. We use a simple Markovian model with two possible environmental states 1 and 2. The probability that the environment changes from state *i* in one generation to the other state in the next generation is *ρ_i→j_*. The total frequency of generations in environmental state 1 is then 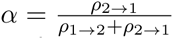.Genotypic fitness of strategies that have a maternal effect depend on the frequency of the two-generation transitions which can be easily calculated. For example, the frequency of generations where there is a transition from environment 1 to 2 is just *ρ*_1→2_*α*.

### 2.2 Fitness of Maternal Effect Strategies

In one extreme, we can consider a maternal effect that allows a mother to choose the distribution of phenotypes that her offspring will have, without any constraints or tradeoffs at any level. In this scenario, the genetic effect is a maternal “strategy” whereby the mother bases the offpsring phenotypes on information that she has about the likely environments her offspring will face. One way for this to occur, for example, is if mothers can alter the developmental trajectory of offspring by adjusting the embryonic environment. This corresponds with an assumption that mothers are free to adjust the fraction of offspring expressing one of two possible phenotypes.

Physiological trade-offs likely limit the scope of maternal effects. For example, the well known size/number trade-off means that allocation of resources to produce larger eggs necessarily limits the number of eggs that can be produced (Smith and Fretwell, 1974; Fischer et al., 2011). We incorporate such trade-offs by assuming that the total number of offspring produced trades-off with offspring phenotype. This could occur because of size-number trade-offs, or because the maternal strategy requires altering maternal physiology or life-history in a way that reduces fecundity. We assume that the total fecundity of females producing a mixed-clutch of offspring is a linear function of her allocation to each phenotype.

We use the geometric mean of reproductive output as the measure of invasion success (Cohen, 1966; Seger and Brockman, 1987). In our case, there is an interaction between pairs of generations so that we must consider not just the frequency of generations of type *i*, but rather the frequency of all 4 possible transitions. Given those frequencies, we can calculate the geometric mean fitness of a genotype based on the reproductive output of the strategy.

Most of the results for the evolution of bet-hedging and maternal effects can be explained by comparing fitness outcomes based on two possible offspring phenotypes. This does not mean that we restrict our attention to two pre-determined phenotypes, but rather consider how maternal effects evolve with two phenotypes at a time followed by mutational input that creates new, but similar, phenotypes.

We call a genotype that produces only a single phenotype of offspring a “pure genetic” strategy since phenotype is determined strictly genetically. If we consider two pure genetic strategies (phenotype 1 and phenotype 2), one will have higher geometric mean fitness and go to fixation. The Log geometric mean fitness of pure genetic strategy 1 and 2 are

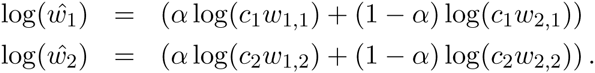

We adopt the parameterization that *c*_2_ = ς_1,2_*c*_1_ and *w_i_*_,2_ = *δ_i,_*_2_*w_i_*,_1_. When considering just two phenotypes we can simplify the notation by always referencing the parameters based on a change from phenotype 1 to phenotype 2 and suppressing the second subscript, i.e. *δi ≡ δi*, 2 and *ς =* ς_1,2_. Assuming that *δ*_1_ < 1, *δ*_2_ *>* 1, we find that

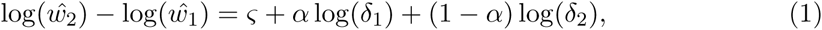

showing that the absolute fecundity and offspring fitness levels are not required to determine which strategy has higher long-term fitness. Because much of the analysis will be done in the *ρ_i→j_* parameter space, it is convenient to find the relationship between *ρ*_1_*_→_*_2_ and *ρ*_2_*_→_*_1_ that favors each of the pure genetic strategies. Solving for the value of *ρ*_2_*_→_*_1_ that causes the fitness difference between strategies to be exactly 0 we find that pure genetic strategy 2 is favored whenever

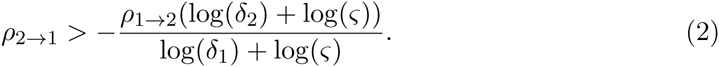

Figure 1 shows the regions of parameter space in which each pure genetic strategy has higher fitness.

**Figure 1:**
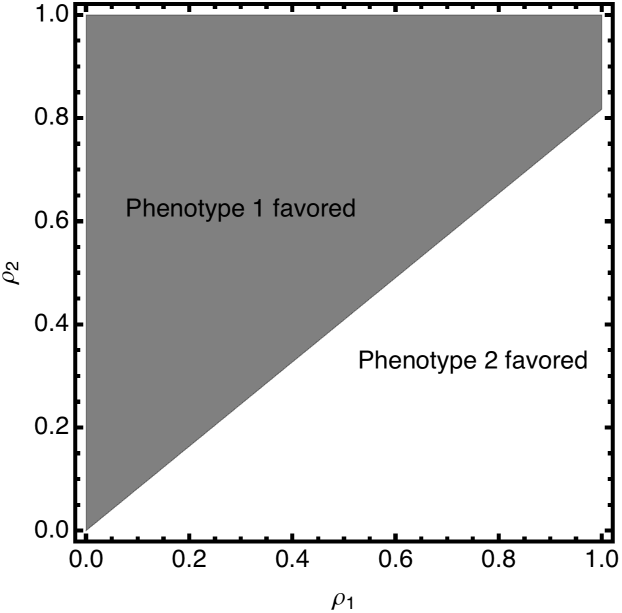
Plots showing regions of parameter space that favor each of the possible pure genetic strategies. Parameters are *δ*_1_ = 0.8, *δ*_2_ = 1.2, ς = 1.0. A line described by inequality 2 divides the parameter space into two regions. Above the line, the frequency of environment 1 is high enough to favor phenotype 1, while below the line phenotype 2 is favored.

#### 2.2.1 Deterministic Maternal Effects

A Deterministic Maternal Effect strategy (DME) is one where the mother determines the offspring phenotype based on her experienced environment. This is a deterministic strategy in that all offspring have the same phenotype. Two possible forms of DME exist, one where mothers produce offspring that have the phenotype adapted to the maternal environment (maintaing DME: mDME), and the other where mothers produce offspring that have the phenotype adapted to the other possible environment (alternating DME: aDME).

To investigate when DME will outcompete pure genetic strategies we have to determine whether or not the fitness of a DME strategy is higher than both possible pure genetic strategies. To do this we first determine which pure genetic strategy has higher fitness and then calculate the difference in log fitness between the DME strategy and the best pure genetic strategy. The log geometric mean fitness of DME strategies that produce offspring with phenotypes 1 or 2 is given by

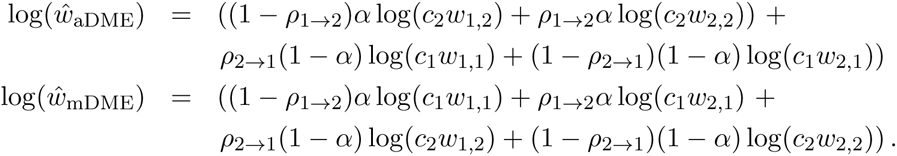

The log geometric mean fitness benefit of the DME strategies can be simplified to

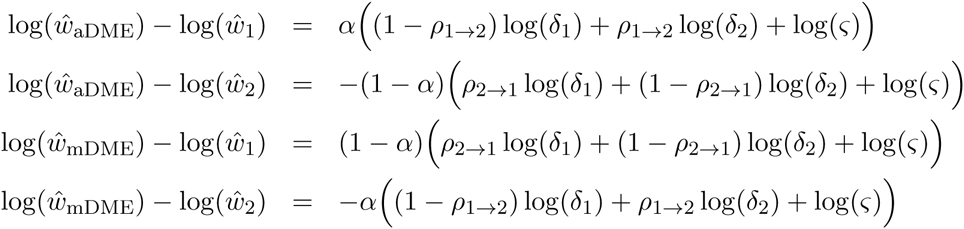

In each case, the advantage of the DME strategy depends simply on a linear function of the log fitness ratio in each environment, the log fecundity cost, and the probability of changing from one environment. To determine which strategy, for a specific set of parameters, represents an ESS we simply test whether the DME strategy can invade the better of the two pure genetic strategies. Figure 4 shows how the ESS depends on the P’s. Assuming that *δ*_1_ < 1 and *δ*_2_ > 1, increasing values of *ρ* make it likely that aDME is the ESS, and decreasing values of *ρ* make it likely that mDME is the ESS. These regions touch at a single point where the line showing that both pure genetic strategies have equal geometric mean fitness crosses the off-diagonal, where *ρ*_2→1_ = 1 − *ρ*_1→2_. This point is defined by

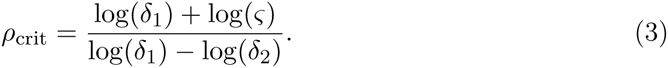

If *ρ*_1→2_ > *ρ*_crit_ and *ρ*_2→1_ > 1 − *ρ*_crit_ then aDME can invade, and if both *ρ*_1→2_ < *ρ*_crit_ and *ρ*_2→1_ < 1 − *ρ*_crit_ then mDME can invade. Otherwise neither mDME or aDME can invade both pure genetic strategies (Figure 2).

**Figure 2:**
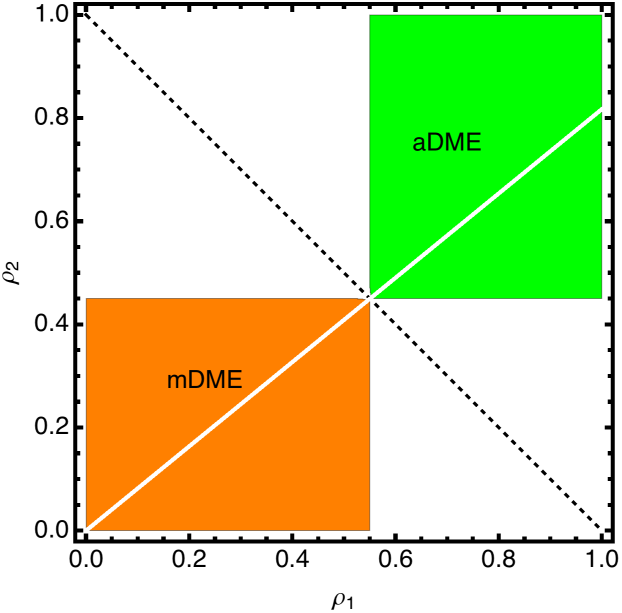
Plots showing regions of parameter space that favor DME. Parameters are *δ*_1_ = 0.8, *δ*_2_ = 1.2, ς = 1.0. The intersection of the line described by inequality 2 and the negative diagonal line defines the critical value *ρ*_crit_. Above and to the right of this point aDME is favored, while below and to the left of this point mDME is favored.

These results imply that DME can only evolve if the environmental sequence is predictable enough, either predictably alternating or predictably remaining constant. Note that when *ρ*_2→1_ = 1 − *ρ*_1→2_ the current environment is completely uninformative in predicting the next generation’s environment (i.e. the mutual information entropy between successive environmental states is 0 (Donaldson-Matasci et al., 2013, 2010)). DME can never evolve if the environment is uninformative, however, for fixed *δ*_1_, *δ*_2_ and ς = 0, then there is always a region where the environments tend to alternate (*ρ*_1→2_ + *ρ*_2→1_ > 1) where aDME is favored, and a region where the environment tends to stay the same (*ρ*_1→2_ + *ρ*_2→1_ < 1) where mDME is favored.

#### 2.2.2 Randomized Maternal Effects

Seger and Brockman define bet-hedging as a reduction in the generation-to-generation variance in reproductive success that increases geometric mean fitness (without increasing the mean of fitness) (Seger and Brockman, 1987), but the term is most commonly associated with one specific mechanism diversification of offspring phenotypes (Kuijper et al., 2014; Simons, 2009; Philippi and Seger, 1989; Crean and Marshall, 2009, e.g. see). We will refer to maternal effects that produce multiple offspring types without using any predictive information as a Randomized Maternal Effects or RME. This is both to avoid existing confusion between bet-hedging and fitness variance reduction, and to emphasize that maternal effects may involve both deterministic and random components.

To determine when RME will invade a population composed of genotypes that code directly for phenotype, we again calculate the geometric mean fitness of the competing strategies. In the case of RME, however, the geometric mean fitness does not depend on the environmental transition probabilities directly, but instead depends on the frequency of the environment 1 ( *α*).

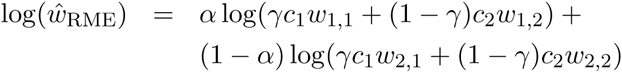

where here *γ* is the probability of producing phenotype 1 and does not vary with the maternal environment. We can now solve for the value of *γ*^*^ that maximizes *ŵ*_RME_. Taking the derivative with respect to *γ* and solving for the critical point gives.

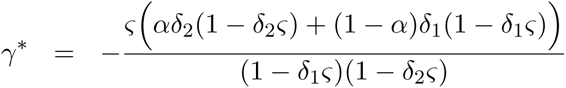

A tedious but straightforward calculation verifies that the second derivative with respect to *γ* is always negative and therefore that W(RME) is maximized at *γ*^*^ if 0 **<** *γ*^*^ < 1. If *γ*^*^ is outside the range of (0,1) then a purely genetic strategy always has higher fitness than RME.Thus, RME can invade the pure genetic strategy if

than RME.Thus, RME can invade the pure genetic strategy if

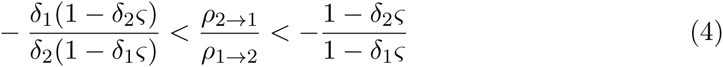

Put simply, if the fitness effect of a change in phenotype is large in both environments (but in opposite direction), then RME is favored for a larger range of *α* values (Figure 3. However, for moderate values of *δ*, the range of *α* that favors RME is fairly restrictive. For example, even with a 2-fold advantage of both phenotypes in their favored environments, only 1/3 of the possible values of a favor the RME strategy.

**Figure 3:**
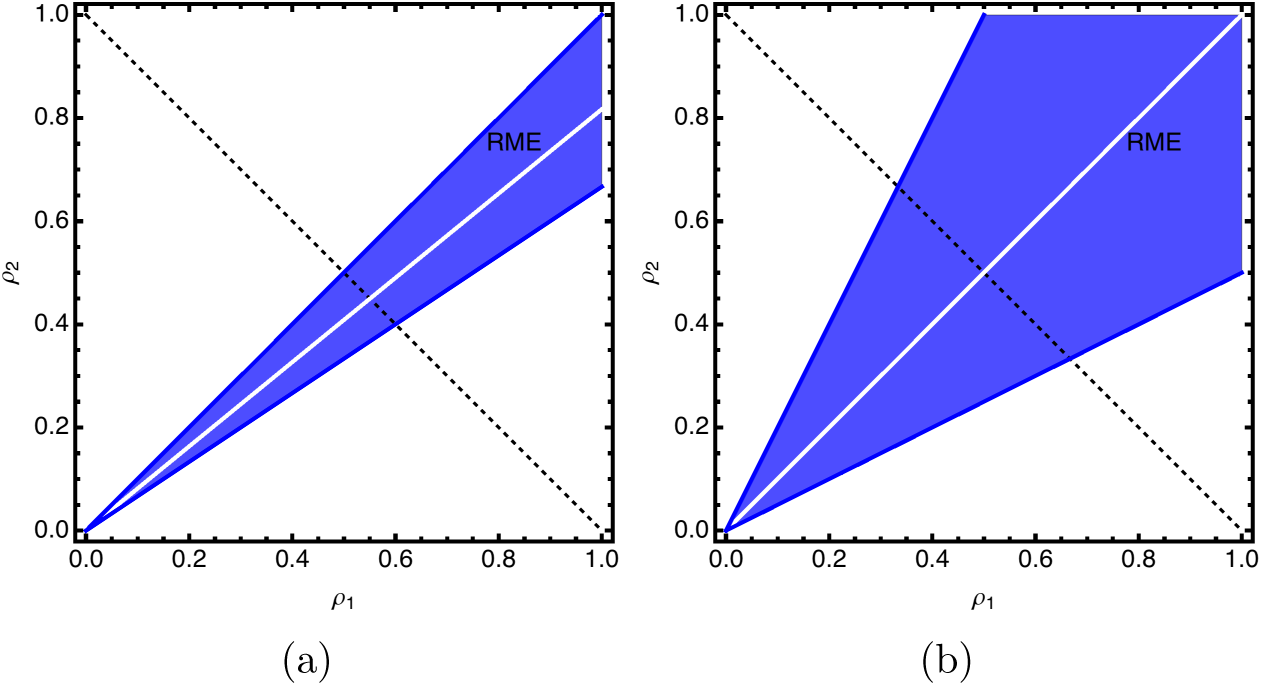
Plots showing regions of parameter space that favor RME. The line described by inequality 2 is shown in white and is always in the center of the cone favoring RME. The width of the region favoring RME depends on the magnitude of the fitness effects. In panel a, parameters are *δ*_1_ = 0.8, *δ*_2_ = 1.2, ς = 0.1. In panel b, parameters are *δ*_1_ = 0.5, *δ*_2_ = 2.0, ς =1.0.

#### 2.2.3 Partial DME

The DME strategies that we have so far considered allow for maternal strategies that use information about the state of the environment to determine offspring phenotype, while RME strategies do not use information about the environment and instead produce offspring phenotypes at random. A more comprehensive strategy, however, is one that uses information about the maternal environment to optimally control the offspring phenotype distribution. This is sometimes called ‘dynamic bet hedging’ or ‘diversified bet-hedging around the norm of reaction’ (Crean and Marshall, 2009; Furness et al., 2015). We refer to this as a hybrid maternal effect, HME. Because a pure genetic strategy, DME and RME are subsets of this strategy, the optimal HME must have higher fitness than all other strategies. Instead of asking when HME is the highest fitness strategy we ask two questions; does the HME strategy collapse onto one of the other strategies and if it does not, what is the magnitude of the fitness advantage in favor of HME?

The HME strategy can be defined by two parameters, γ*_i_*_,1_, which determine the probability of producing offspring of phenotype 1 when the mother experiences environments *i*. For simplicity, we define γ*_i_* = γ*_i_*_,1_. The log geometric mean fitness benefit of the HME strategy can be simplified to

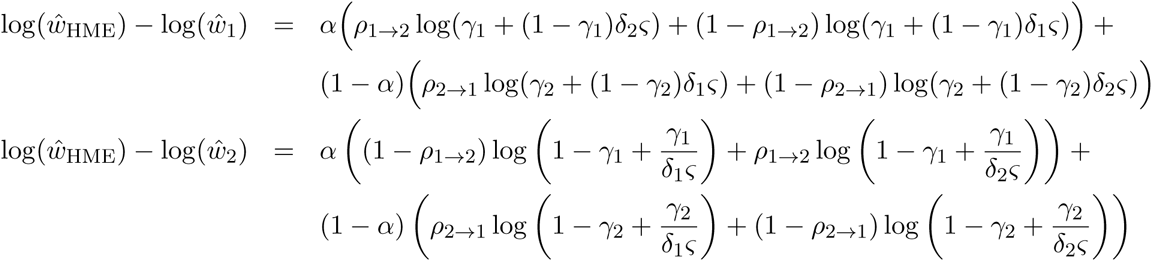

We can find the values of γ_1_, γ_2_ that maximize the fitness of the HME strategy by taking the derivative and solving for the critical point. Again, only if the critical point has 0 < γ*_i_* < 1 will this represent the optimum value. We find

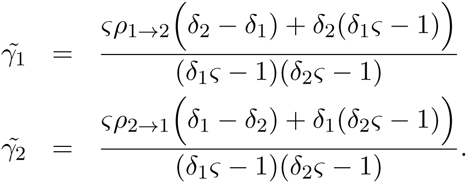

While HME always has fitness greater than or equal to the other maternal effect strategies, the optimal γ*_i_* may be 0 or 1, indicating that HME strategy collapses to a DME strategy. In fact, only a small segment of the (*ρ*_1→2_, *ρ*_2→1_) parameter space favors HME in this stricter sense. By solving for the values of (*ρ*_1→2_, *ρ*_2→1_) that lead to 0 < γ*_i_* < 1 gives

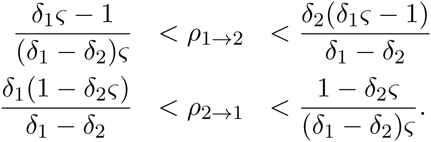

As shown in Figure 4, the area where bet-hedging is strictly favored in both parental environments is represented by a square whose corners are positioned at the intersection of negative diagonal line and the line showing the region where RME is strictly better than a pure genetic strategy (i.e. the lines defined by inequality 4).

**Figure 4:**
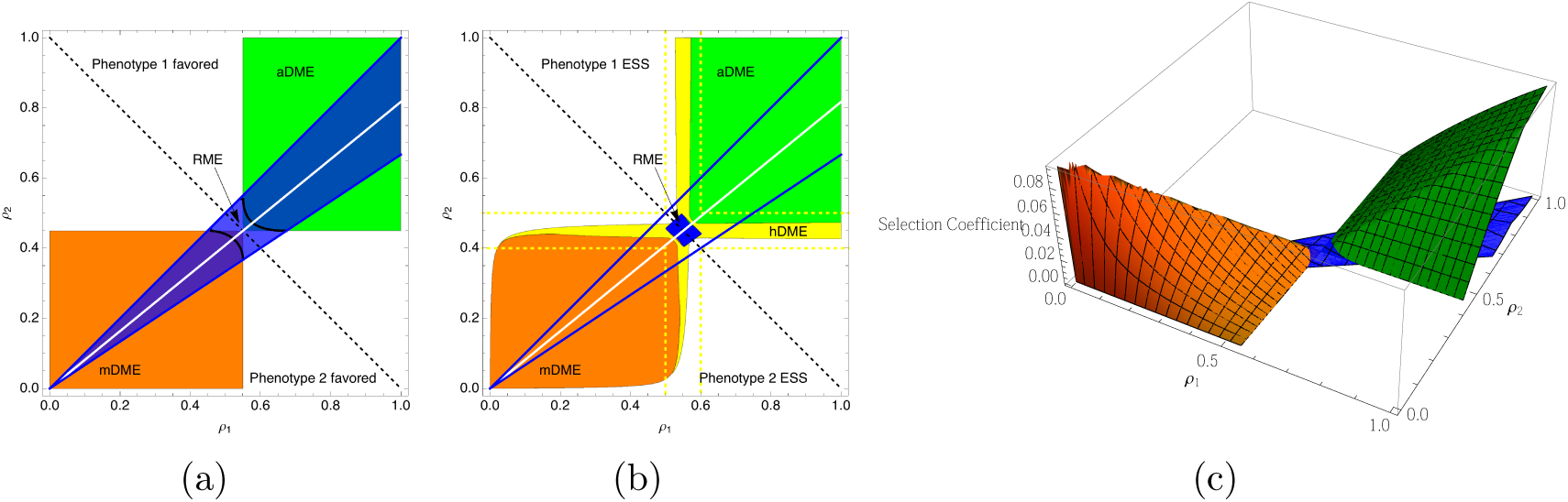
Plots showing regions where different types of maternal effect can invade the single-phenotype strategy. For both panels, *δ*_1_ = 0.8, *δ*_2_ = 1.2, *ς* = 1.0. The dashed line on the diagonal is for reference only. The solid white line represents the boundary between regions where the two pure genetic strategies have equal fitness. The point where the dashed line and the solid white line intersect define the rectangular regions that favor the DME strategies. Panel (a) shows the regions that allow mDME, aDME, and RME to invade the pure genetic strategies. The area where the blue shading overlaps either the orange or green represent parameters that would allow both RME and DME to invade against a pure genetic strategy. However, RME has higher fitness than DME only between the two black curves. Panel (b) shows regions of parameter space that favor different maternal effects strategies if the selection coefficient is larger than 10^−3^. Again, regions between the two blue curves allow for RME to replace pure genetic strategies, but only in the blue area does RME invade DME. The HME strategy is predicted to have the highest fitness in the region defined by the dashed yellow lines, however the selection coefficient in favor of HME is larger than 10**^-^**^3^ in the region shaded with yellow. Panel (c) shows the fitness surface for aDME, mDME and RME.

### 2.3 ESS maternal effect strategies

In this section we consider competition between maternal effect strategies. For example, two discrete phenotypes may be available (i.e. shade and sun leaves), and we consider competition between alternative maternal effect strategies that change when the two phenotypes are produced, but do not alter the phenotypes themselves.

The expressions for the fitness benefit of the different maternal effect strategies can be used to determine which strategy has the highest fitness for a specific set of parameters. Although genetic and ecological interactions could cause frequency-dependent effects, no frequency-dependence emerges from the basic model. This means that the outcome of selection can be predicted based on the genotypic fitness values directly. Thus we can consider a fixed set of phenotypic fitness values and determine which regimes of environmental change favor specific maternal effect strategies over pure genetic strategies.

First considering only RME and DME, we find that in most of the region where DME (either aDME or mDME) has higher fitness than the pure genetic strategy, DME also has higher fitness than RME. When the parental environment provides little information about the offspring environment, then RME has higher fitness than DME. As the level of environmental predictability goes up, fitness of the DME strategies increase while the fitness of RME strategies does not (it depends only on *α* (figure 4 panel a). Note that RME and DME must have equal fitness at the point where they also have equal fitness with the pure genetic strategy, i.e. at the points where the green and blue or orange regions intersect. Since increasing the difference in fitness between the phenotypes in the two environments increasing |log(*δ_i_*) |) leads to a wider cone described by the blue region, increased fitness effects also result in a larger area where RME is favored.

We also need to consider HME, but since HME always has fitness greater than or equal to the other strategies it makes more sense to ask when a mutation that causes a change in maternal effect strategy would be likely to spread in finite population. To do this we simply assume that there is a selection coefficient threshold, *s*_min_, below which mutations are unlikely to become fixed (i.e. because population size is finite and drift is expected to dominate at low values of *s*). We consider the sequential introduction of RME, DME and HME (the alternative sequence of DME, RME, HME yields similar results).

An example shown in figure 4 b serves to illustrate the general results. The slope of the white line depends on the fitness parameters and the fecundity cost, *ς*, and remains in the upper right quarter plane so long as *ς* is not too different from 1. The conditions that favor RME are restricted to a small region around the negative diagonal, around which the HME strategies have the highest fitness. HME is never under very strong selection when compared with RME and DME, but has the largest effect near the intersection of the RME region and the DME regions. This is evidenced by the fact that the region where selection in favor of HME is larger than *s*_crit_ is hugs the contours of the regions favoring DME. Although there is complete symmetry in the boundaries of the regions where aDME and mDME have higher fitness than the pure genetic strategy, panel b shows that there is a quantitative difference in these effects as shown by the smaller area of the orange region when compared with the green region.

The size of the regions favoring DME, RME, and HME depend on the parameters determining environments specific fitness and phenotype dependent fecundity. Increases in the environment-specific effects make the regions supporting RME and HME larger, at the expense of the size of the regions favoring DME (figure 5). Changes in the environment-specific selection coefficients can shift the regions that favor DME, but RME is still favored in environmental regimes that carry little information about the future.

**Figure 5:**
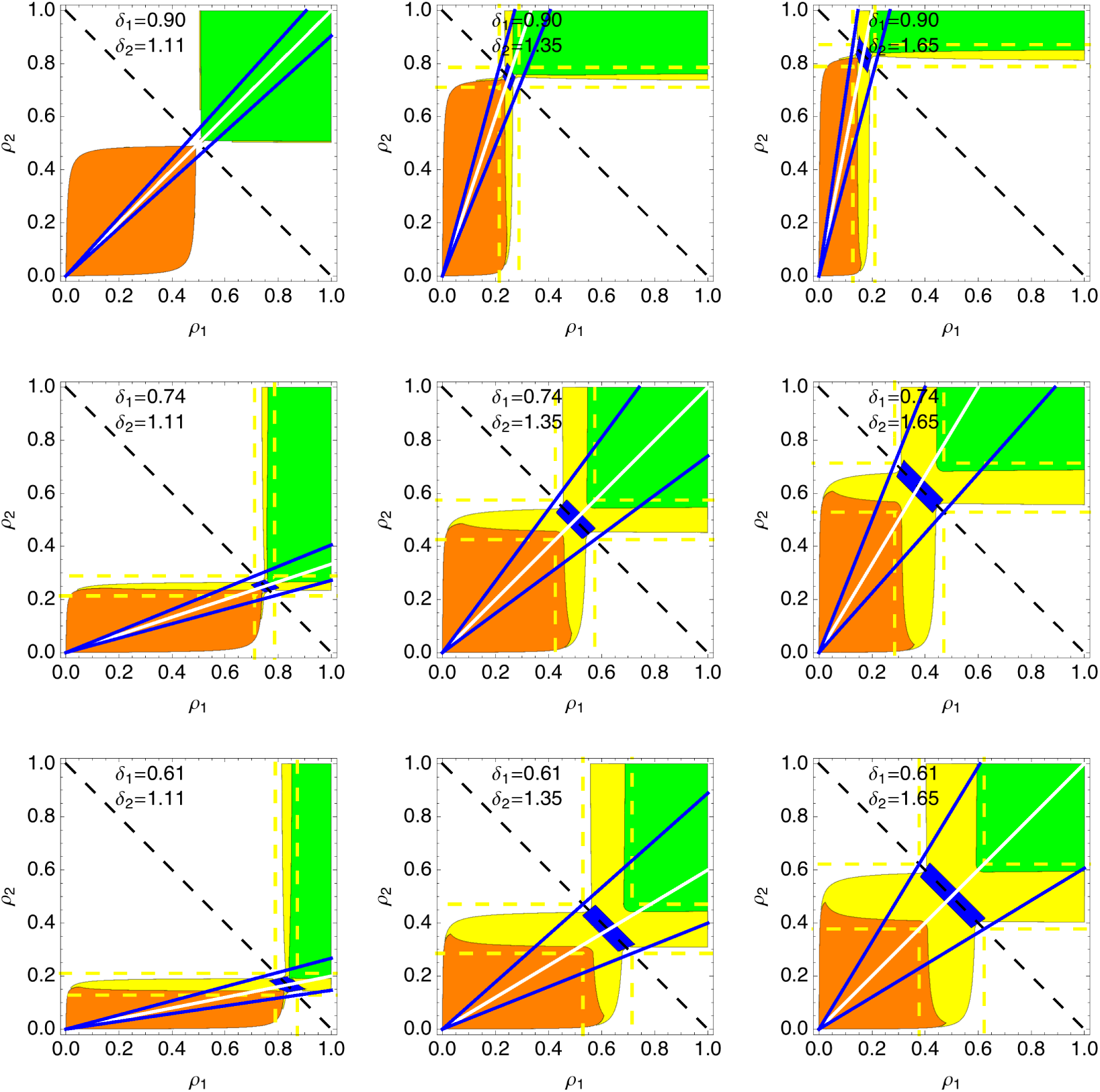
The effect of the within-environment selection strength on the regions where alternative forms of maternal effect are favored. aDME is favored in the green regions, mDME in the orange, RME in the blue, and HME in the yellow. The critical selective advantage is set at *s*_min_ = 10**^-^**^3^ and both phenotypes have the same fecundity effect, *ς* = 1. The values of *δ*_1_, *δ*_2_ were systematically varied from *e*^−(.1+.2*^*^j^*) and *e*^.1+.2^\**^j^*, *j* from 1 to 3, respectively.

Changes in the fecundity cost (*ς*) also shift the regions favoring maternal effects, but maintains the same relative range of *α* that allow RME (figure 6). As expected, when one phenotype is favored in terms of maternal fecundity, mothers benefit by producing the easier to produce phenotype. If this effect is large enough, then the pure genetic strategy to produce the phenotype associated with higher fecundity is the ESS regardless of the environmental regime. The region supporting pure RME remains quite small, regardless of the change in *ς*.

**Figure 6:**
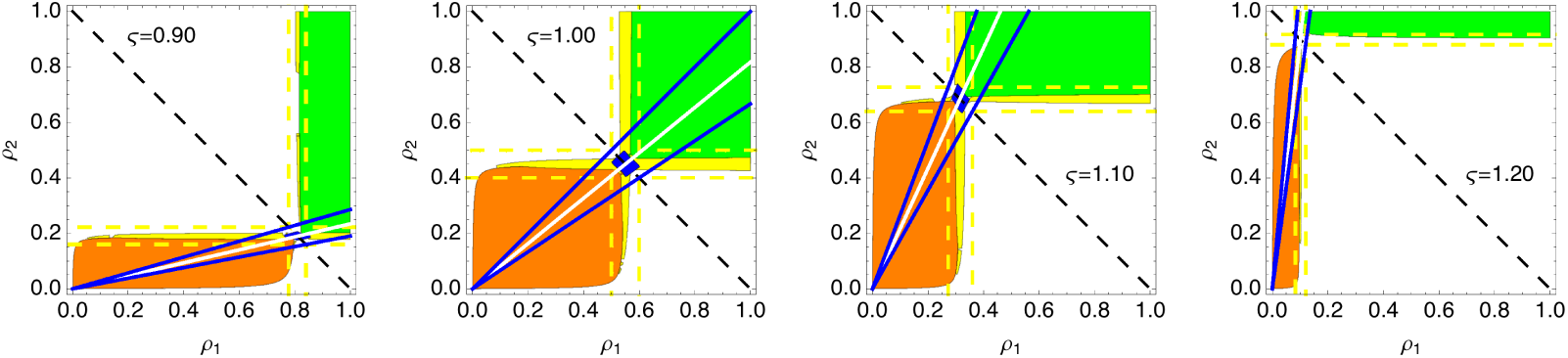
Effect of differential fecundity of the two phenotypes on the regions where alternative forms of maternal effect are favored. For all panels, aDME is favored in the green regions, mDME in the orange, RME in the blue, and HME in the yellow. The parameter values are *δ*_1_ = 0.8, *δ*_2_ = 1.2, and *s*_min_ = 10^−3^. *ς* was systematically varied from 0.9 through 1.2. Lower and higher values of *ς* create very strong selection in favor of a single phenotype meaning that one of the pure genetic strategies is the ESS.

**Figure 7:**
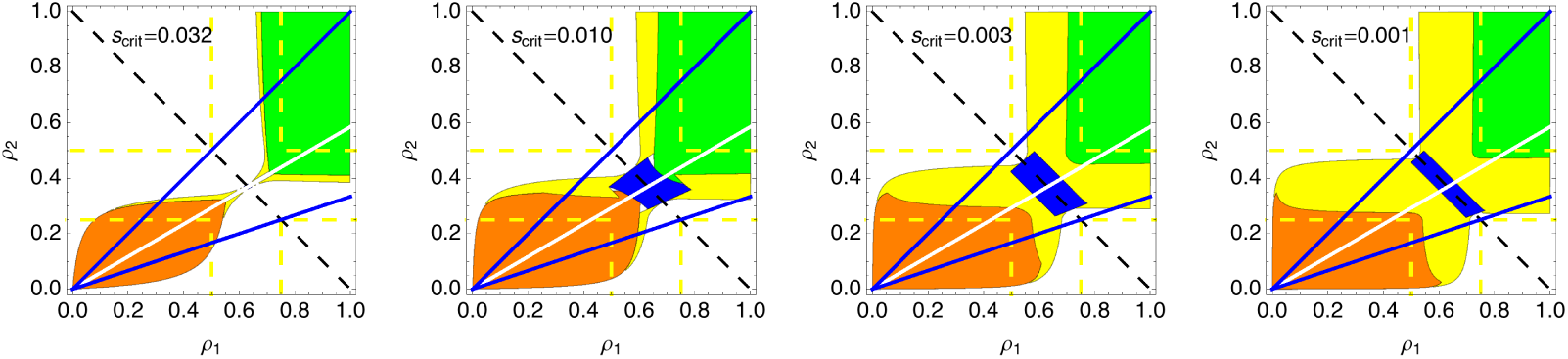
Effect of the threshold value of the selection coefficient, *s*_crit_. For all plots, aDME is favored in the green regions, mDME in the orange, RME in the blue, and HME in the yellow. When *s*_crit_ is high the region that supports RME is absent. Larger values of *s*_crit_ give RME more scope to evolve, but also increase the opportunity for HME to replacer RME. As the value of *s*_crit_ increases, the region supporting HME increases to fill the region within the dashed yellow lines.

**Figure 8:**
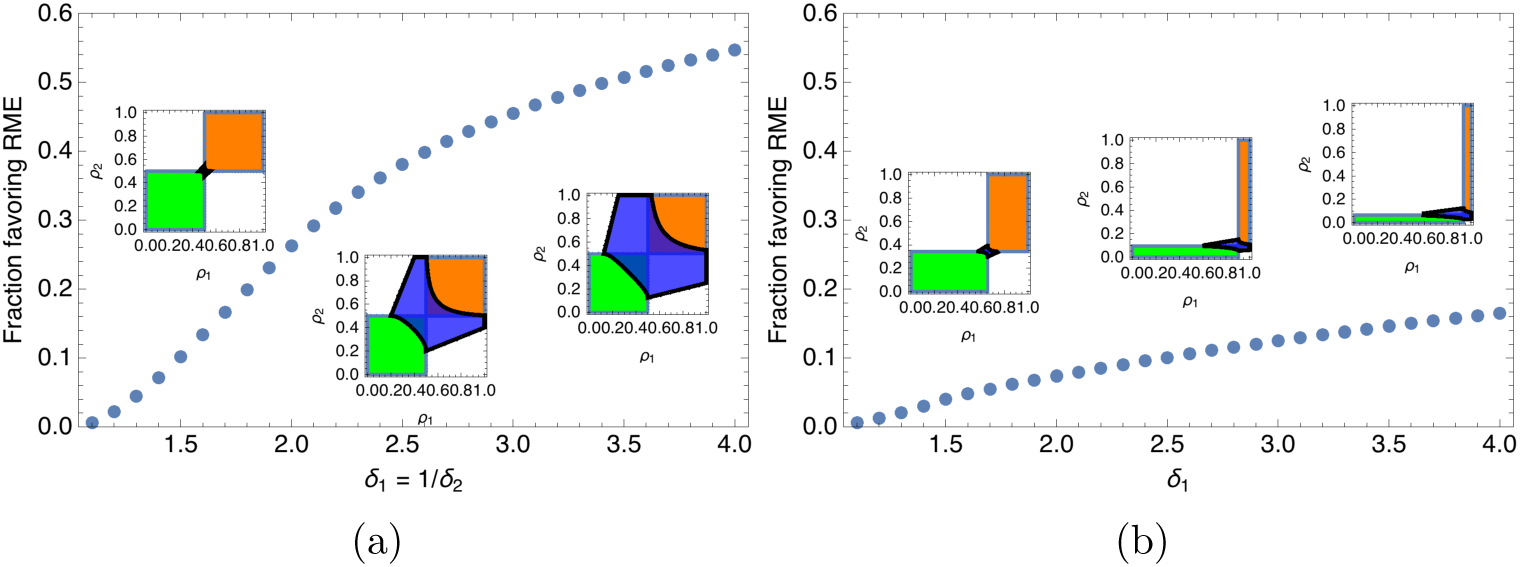
Fraction of parameter space that favors RME as compared with DME. In both panels, the vertical axis is the are of parameter space favoring the RME divided by the area of parameter space where either RME or DME is favored over a single phenotype. There is no fecundity advantage, so *ς* = 1. Panel (a) shows the response when allele 2 is disfavored in environment 1 by the same amount as it is favored in environment 2. Therefore *δ*_1_ = 1/*δ*_2_. The magnitude of the fitness difference is varied from *δ*_1_ = 1.1 to *δ*_1_ = 4.0. Inset diagrams show the area where DME and RME are favored. Panel (b) shows the effect of asymmetric environment-specific fitness. In this panel, *δ*_2_ = 0.91 and *δ*_1_ is varied between 1.1 and 4.

While changes in *s*_crit_ alter the range of parameters that support all types of maternal effect, it has a much larger effect on HME and RME (figure 6). This is because the fitness advantage of DME is generally much larger than for RME. Increasing *s*_crit_ first increases that area that supports pure RME, but as *s*_crit_ gets larger HME tends to replace RME.

## 3 Joint evolution of phenotype and maternal effects

In the previous sections, we considered how mutations that produce a novel maternal effect strategy would fare. However, both the phenotypes and the strategic deployment of those phenotypes are expected to evolve. In this section we consider how a population can evolve both in terms of the phenotypes produced as well as in terms of the strategies controlling phenotype production. We first consider mutations that introduce a novel phenotype into an ancestral population that produces a single phenotype in all situations. We then examine the joint invasion of a maternal effect along with a change in phenotype. We then present some numerical examples and simulations based on a model of phenotypic trade-offs.

### 3.1 Invasion of alternative phenotypes

In this section we turn to evolutionary change in the phenotypes that are available to be produced by the maternal effect strategy. This allows us to consider mutations that cause small changes in the phenotypes from an ancestral state where only a single phenotype is produced. These results are useful in their own right, but also pave the way to considering the joint evolution of the phenotypes and the maternal effect strategy in the next section.

We start by considering a population that produces a single phenotype in all offspring. We then ask how a mutant that has a specific maternal effect strategy that produces the ancestral phenotype and an alternative phenotype. This is like introducing a mutant that has a joint effect both on the strategy of phenotype production and on the phenotypes that are produced. The idea is that one of the phenotypes is the original (ancestral) phenotype and the other only differs from the ancestral phenotype by a small amount.

When the values of *ρ*_1→2_, *ρ*_2→1_ are in a region that provides little information about the future environment, then only RME can invade. However, the values of *δ*_1_, *δ*_2_ must be finely balanced. If the benefit in environment 1 is too large, then the purely genetic strategy using the novel phenotype would invade and replace the ancestral phenotype. Likewise, if the fitness cost in environment 2 is too large then the bet-hedging strategy will not invade at all. In fact, the curves defining the region where RME can invade approach each other faster than linearly as the difference from the ancestral phenotype becomes small. This means that only very specific phenotypic trade-offs can possibly support invasion of RME. Figure 9 shows that RME only invades for a narrow range of fitness parameters and, when *s*_crit_ > 0 the change in phenotype must be above a threshold for invasion.

**Figure 9:**
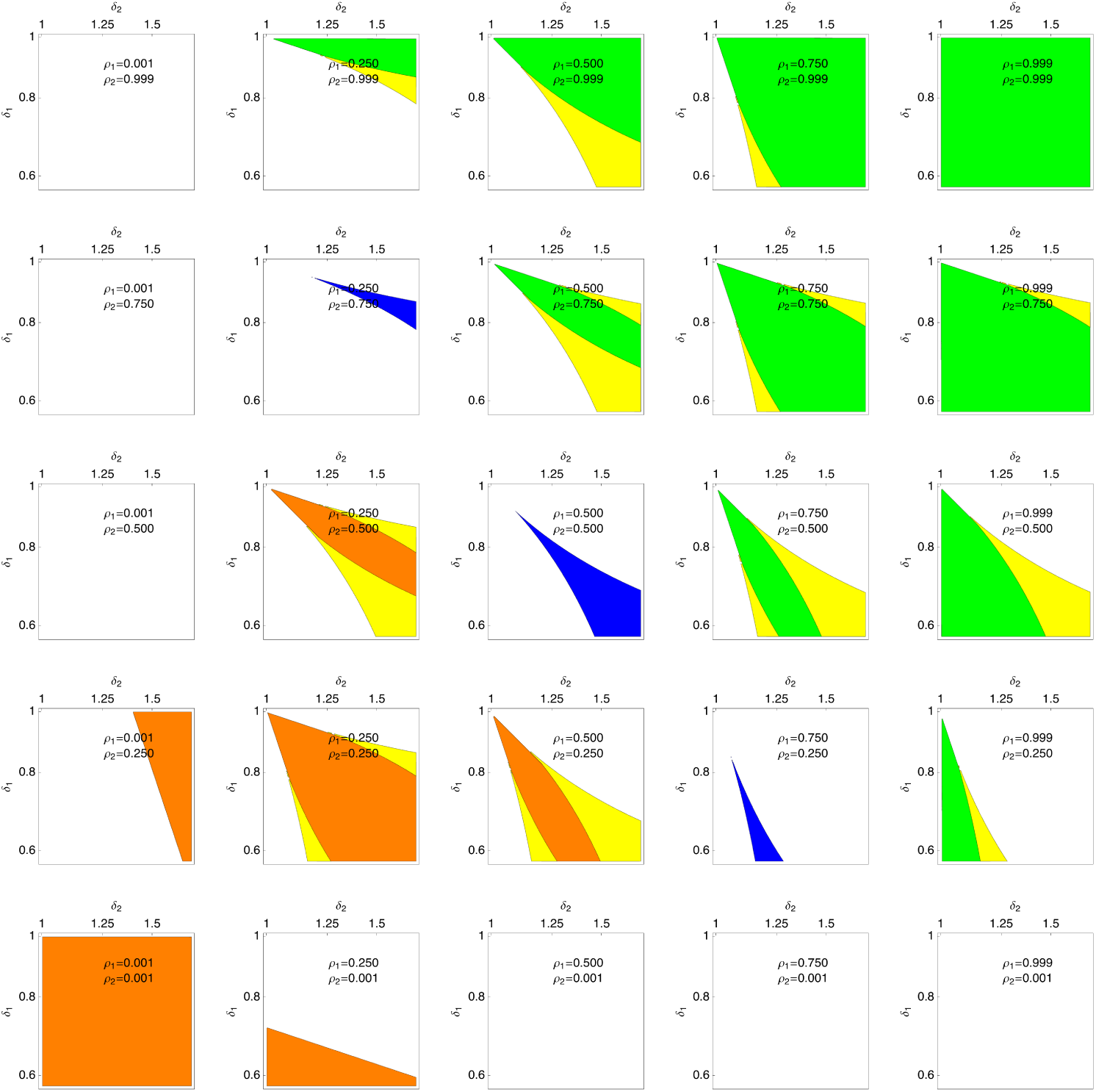
Invasion of a novel phenotype using fixed maternal effect strategies. The colored regions show parameters for a mutant that produces a novel phenotype that decreases fitness in condition 1 and increases fitness in condition 2. Note that a phenotype that is the same as the ancestral phenotype would be represented by a point in the top left corner of the plot. The axes are on a log scale. For all panels *s*_crit_ = 0.001 and *ς* = 1. For all plots, aDME is favored in the green regions, mDME in the orange, RME in the blue, and HME in the yellow. For values of *ρ*_1→2,_ *ρ*_2→1_ that contain little information about the environment (i.e. *ρ*_2→1_ = 1 − *ρ*_1→2_) RME can invade, but the region of invasion becomes narrow as the mutant effect is small. For larger values of *ρ*, aDME can invade, and HME is only favored in a narrow slice of parameter space. For lower values of *ρ*, mDME can invade, but the regions of invasion are smaller than for aDME.

When the maternal environment does contain significant information about the offspring environment, then DME can invade for a range of values of *δ*_1_, *δ*_2_. So long as *ρ*_1→2_ ≠ 1 − *ρ*_2→1_, the region allowing invasion DME is bounded by two curves that have different slopes, even near the ancestral phenotype. This means that there are always a range of phenotypic trade-offs that would allow invasion of DME. Both aDME and mDME have similar ranges of invasion when the rates of environmental change are symmetric (i.e.*ρ*_1→2_ = *ρ*_2→1_), but mDME is substantially more limited for asymmetric environmental transition rates (Figure 9)

### 3.2 Joint evolution of maternal effect and phenotypes

In this section we consider an ancestral state of a purely genetic strategy that produces one phenotype in all conditions. We then allow mutations to introduce variance in the phenotype, and allow the population to move to a stable attractor, the single-phenotype ESS. At this ESS, the first order selection coefficients for changes in phenotype are, by definition, 0. Once the population has reached the single-phenotype ESS we look at mutations that have small effects on both the phenotypes produced and the frequency at which the phenotypes are produced. These can be mutations that produce RME or mutations that produce HME.

First we derive the conditions for the spread of mutations that cause HME or RME. In this context HME involves a small frequency of offspring carrying a new phenotype to be produced after the mother experiences one environment (but not the other). Further, the new phenotype is assumed to be quantitatively similar to the single-phenotype ESS phenotype. This notion of the phenotype being produced only infrequently and being a similar phenotype to the ancestral phenotype is analogous to looking at the second order effects of changes in phenotype. We show that once the population reaches the single-phenotype ESS, mutations that introduce HME are favored so long as*ρ*_1→2_ ≠ 1 − *ρ*_2→1_. In other words, HME can evolve as long as the parental environment provides some information about the offspring environment.

Likewise, we can explore selection on mutations that introduce RME near to the single-phenotype ESS. Here we mean a mutation that causes a new phenotype to be produced infrequently irrespective of the parental environment. Again, the new phenotype is assumed to be quantitatively close to the single-phentoype ESS. In contrast to the HME result, second order selection in favor of mutations that causes RME is always 0.

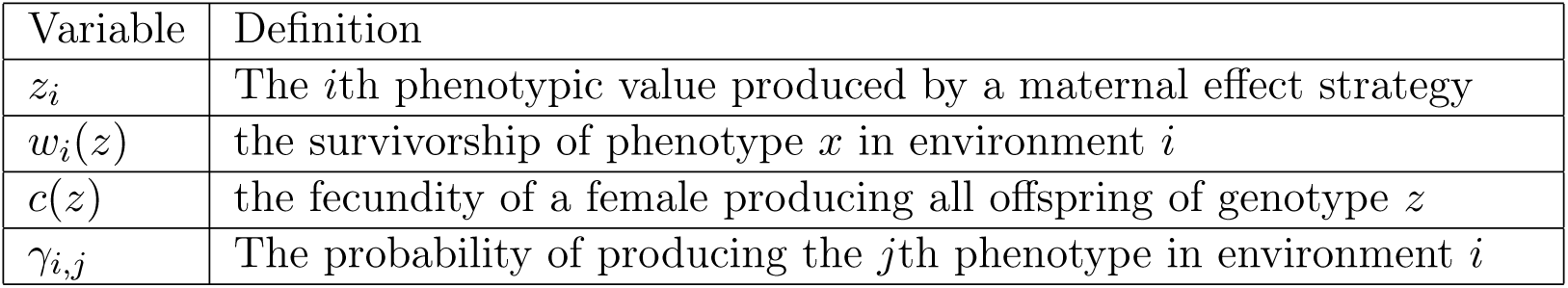

| Variable | Definition |
| --- | --- |
| $z_i$ | The $i$th phenotypic value produced by a maternal effect strategy |
| $w_i(z)$ | the survivorship of phenotype $x$ in environment $i$ |
| $c(z)$ | the fecundity of a female producing all offspring of genotype $z$ |
| $\gamma_{i,j}$ | The probability of producing the $j$th phenotype in environment $i$ |

In the absence of any type of maternal effect or phenotypic randomization, we expect that the population will evolve to an ESS where all individuals have a single phenotype that balances the selective pressures from both environments. We assume that there is a single dimension that describes the phenotype, that the two environments favor different phenotypes via fitness functions that are monotonic in the region of interest, and that female fecundity depends on offspring phenotype alone. Fecundity is assumed to be a monotonic function of phenotype. Without loss of generality we assume that

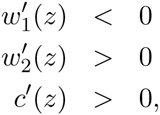

such that environment 1 favors lower values of *z*, environment 2 favors higher values of *z*, and fecundity is highest for low values of *z*. This means that any hybrid maternal effect would involve increasing the value of *z* in anticipation of offspring experiencing environment 2 and decreasing the value of *z* in anticipation of offspring experiencing environment 1. We write the general geometric mean fitness function as a function of four variables, *ŵ*(*z*_1_ *z*_2_, *γ*_1,1_, *γ*_2,1_). Note that if *γ*_1,1_ = *γ*_2,1_ = 1 then only phenotype *z*_1_ is expressed. We can define

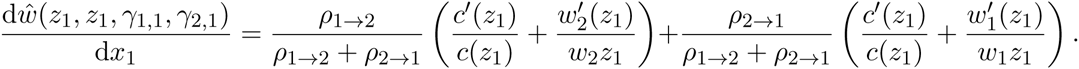

Solving for the value of *z*_1_ = *z*^*^ that makes the derivative of fitness zero gives the single-phenotype ESS,

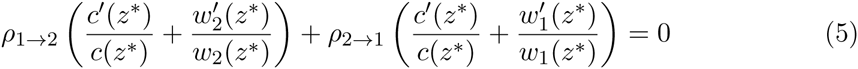

Now we would like to determine the spread of mutants that alter the phenotypes *z*_1_, *z*_2_ as well as the environment-specific probability of producing each of those phenotypes. Our main simplifying assumption here is that mutations will cause only a small change in the phenotype. In contrast, we are able to explore both infinitesimal and large changes in *γ*.

We assume the population has already evolved to the single-phenotype ESS and that *γ*_1_ = *γ*_2_ = 1. This represents a genotype where only a single phenotype is produced, and avoids artificial conditions such as genotypes that code for an phenotype that is never produced. If we write the fitness function as *ŵ*(*z*_1_ *z*_2_, *γ*_1,1_, *γ*_2,1_) then the single-phenotype ESS is characterized as *ŵ*(*z*^*^, *z*^*^, 1,1). Altering either *γ* has no effect on fitness and altering the value of the second phenotype (i.e. *z*_2_) has no effect on fitness because these changes do not alter phenotype in any situations. This means that

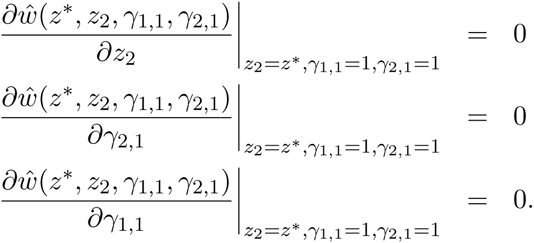

Simply put, mutations that do not alter the phenotype have no effect on fitness. This is, to some extent, due to the choice to parameterize the phenotypic values *z*_1_ and *z*_2_ as separate parameters from *γ*_1,1_ and *γ*_2,1_.

Invasion of a mutant that creates either pure bet-hedging or a hybrid maternal effect can be worked out by considering mutant strategies near the single-phenotype ESS that alter both *z*_2_ and the *γ*’s. For bet-hedging, this is a mutation that alters both *γ*_1,1_ and *γ*_2,1_ by the same amount, whereas a hybrid maternal effect decreases only one *γ* and takes advantage of the correlation between the parental environment and the offspring environment.

Invasion of the mutant depends on the appropriate mixed partial derivative. For bet hedging to invade, we require 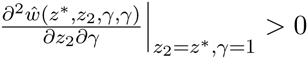. However, we find that

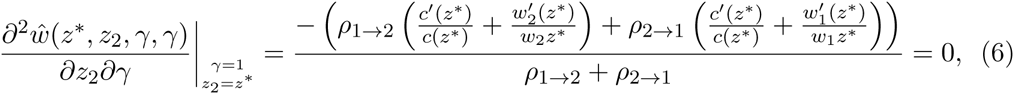

which is zero because of the condition at the single-phenotype ESS. Thus, bet-hedging strategies that have small changes in the phenotype do not, to a first order approximation, cause an increase in fitness. This is true regardless of the fitness functions or environmental frequencies. This approach can easily be extended to third order effects, where it can be seen that positive selection for RME is sometimes, but not always, present.

For hybrid strategies, only one of the *γ* values is altered, and this is done in a way that depends on the fitness values of the phenotypes. Obviously, mutations could arise that cause the altered phenotype to be produced at inappropriate times, and these would be selected against. The appropriate condition for invasion of a mutation with a small effect on the HME, when an increase over *z*^*^ is favored if the current environment is 1, is 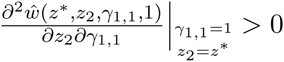. We find that

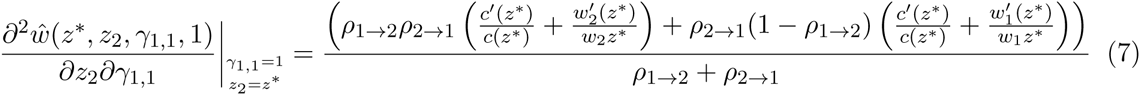

By comparing this expression and the expression for the single-phenotype equilibrium (equation 5) we note that the first term is multiplied by *ρ*_2→1_ while the second by 1− *ρ*_1→2_. By assumption 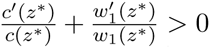 so that whenever *ρ*_2→1_ > 1 − *ρ*_1→2_ the maternal effect invades. This is because the fitness derivative taken implies that *γ* is reduced when the mother faces environment 1, and *ρ*_2→1_ > 1 − *ρ*_1→2_ implies that when mothers experience environment 1, their offspring are more likely to experience environment 2 than if the mother herself experienced environment 2. In general, *ρ*_2→1_ > 1 − *ρ*_1→2_ indicates that the environments have a tendency to alternate, whereas *ρ*_2→1_ < 1 − *ρ*_1→2_ indicates that the environments have a tendency to remain the same over multiple generations. A similar calculation shows that 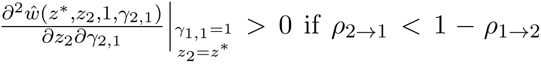, implying that maintaining maternal effects can evolve when environments tend to remain the same. This result shows that regardless of the specific details of the survivorship and fecundity functions, HME can always invade a single-phenotype ESS.

Other possibilities include more extreme types of maternal effect strategies, such as an RME strategy that produces the optimal *γ* or DME strategies that have *γ*_1,1_ = 1, *γ*_2,1_ = 0. Similar calculations can be made and we find that whenever mutations cause small effects in phenotypes but produce those phenotypes in an environment-independent way (i.e. bet-hedging), then the second order fitness effects are 0. In contrast, even complete DME shows a positive second order fitness effect.

### 3.3 Joint evolution of the maternal effect strategy and offspring phenotypes under a phenotypic trade-off

In this section we develop a simple trade-off model for phenotypes and show how the trade-off curve allows for the evolution of maternal effects. As the results from the previous sections show, we find conditions where DME evolve by small mutational steps from ancestral populations that lack maternal effects.

We use, as an example, a model where the fitness in each type of environment is a Gaussian function of the difference between the phenotype and the environment-specific phenotypic optimum. Thus environment specific fitness is defined as

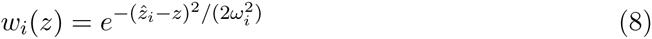

where *z* is the phenotypic trait value, ẑ*_i_* is the optimum phenotype in environment *i*, and *ω_i_* is inversely related to the strength of selection in environment *i*. For simplicity, the environment specific optima are set at *ẑ*_1_ **=**0 and *ẑ*_2_ = 1 and we assume that *w* = *w*_i_. It has been shown by Bull using a continuous trait version of this model that selection for phenotypic diversification depends on *ω* being small relative to the magnitude of the fluctuations in the phenotypic optima (i.e. *ẑ*_2_ − *ẑ*_1_) (Bull, 1987). Fecundity effects can also be tied to the phenotypic trait value by assuming that fecundity is a continuos function of *z*. In the absence of fecundity effects, the environment-specific fitness surface is defined by one parameter, *ω* and overall fitness surface depends on *ω*, *ρ*_1→2_, and *ρ*_2→1_.

In the previous section we showed mutations inducing DME can always invade the single-phenotype optimum, and that conditions for the invasion of RME were more restrictive. We showed for general fitness functions that selection for RME is weaker than selection for DME, because the second order effects are always zero. (As a side note, it is also the case for Bull’s result that invasion of increased phenotypic variance is a higherorder effect.) Given the Gaussian model assumed here, we can calculate whether or note RME can invade at all by examining the mixed partial derivatives of fitness near the single-phenotype optima. Taking the partial derivative of the log of the general fitness function gives

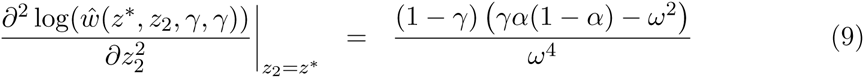

where here *γ* is probability of producing phenotype *z*^*^ and is independent of the maternal environment. If *γ* = 1 then only the phenotype *z*^*^ is produced. For any value of *γ* < 1 the expression will be positive if *ω*^2^ is sufficiently small. Mutants that have *γ* near 1 can invade so long as *ω*^2^ < *α*(1 − *α*). The same result is found if we assume that the RME mutant produces two phenotypes equally spaced on either side of the single-phenotype ESS. As in Bull’s classic result, we find that the fitness function must be sufficiently steep for phenotypic variation to be selected (Bull, 1987). When *ω* is too high to allow invasion of RME, then the ESS must involve DME.

Under the assumption that successive mutations only alter the phenotypes produced by small amounts, we can work out the long-term evolutionary behavior by examining the fitness gradient. Our general results guarantee that near the signle-phenotype ESS, the gradient is steeper for mutants inducing a form of DME than for mutants inducing RME. As successive mutations appear and spread in the population, the common phenotypes in the population get farther from the single-phenotype ESS, making our approximations invalid. Further evolution will occur, and if RME mutants become more fit than the resident DME strategy, for the same phenotype parameters, then we expect that RME will spread and may be the ESS. Based on these ideas, the evolutionary outcome can be divided into 4 types of outcome: (1) RME cannot invade and the ESS involved some form of DME, (2) RME could invade in the absence of DME, however DME represents the global fitness maximum and RME is excluded (3) RME could invade in the absence of DME, and RME represents the global fitness maximum. Even in this case, DME may be an ESS because RME mutants near the optimum DME phenotypes have lower fitness (4) RME can invade and has higher fitness than DME for most parameters; RME is the ESS and can evolve smoothly.

Examples of the 4 general types of evolutionary dynamics are shown in figure 10 and the supplemental *Mathematica* notebook can be used to explore the parameter space further. Panel (a) in figure 10 shows an example where the curvature of the fitness surface precludes the evolution of RME. In this example, the environment-specific fitness functions are relatively shallow. This can also be expressed in terms of a Levins’ fitness set, where the possible values of fitness are plotted against each other (Levins, 1962). The curvature of the fitness set is concave. Our general result for invasion of DME can be shown in terms of the fitness effects that are required for DME to invade, and on top of that we plot (on a log scale) the fitness set. Because the fitness set passes through the green region we know that mutants that alter one phenotype but have DME will invade. We can also visualize the effect of mutations that alter both phenotypes by plotting the regions where DME can invade the single-phenotype ESS along with the contours of the fitness function. We can see that mutations that push one *z* value higher and the other lower will invade and that the joint maternal effect and phenotypic value ESS occurs at *z*_1_ = .75, *z*_2_ = .25.

**Figure 10:**
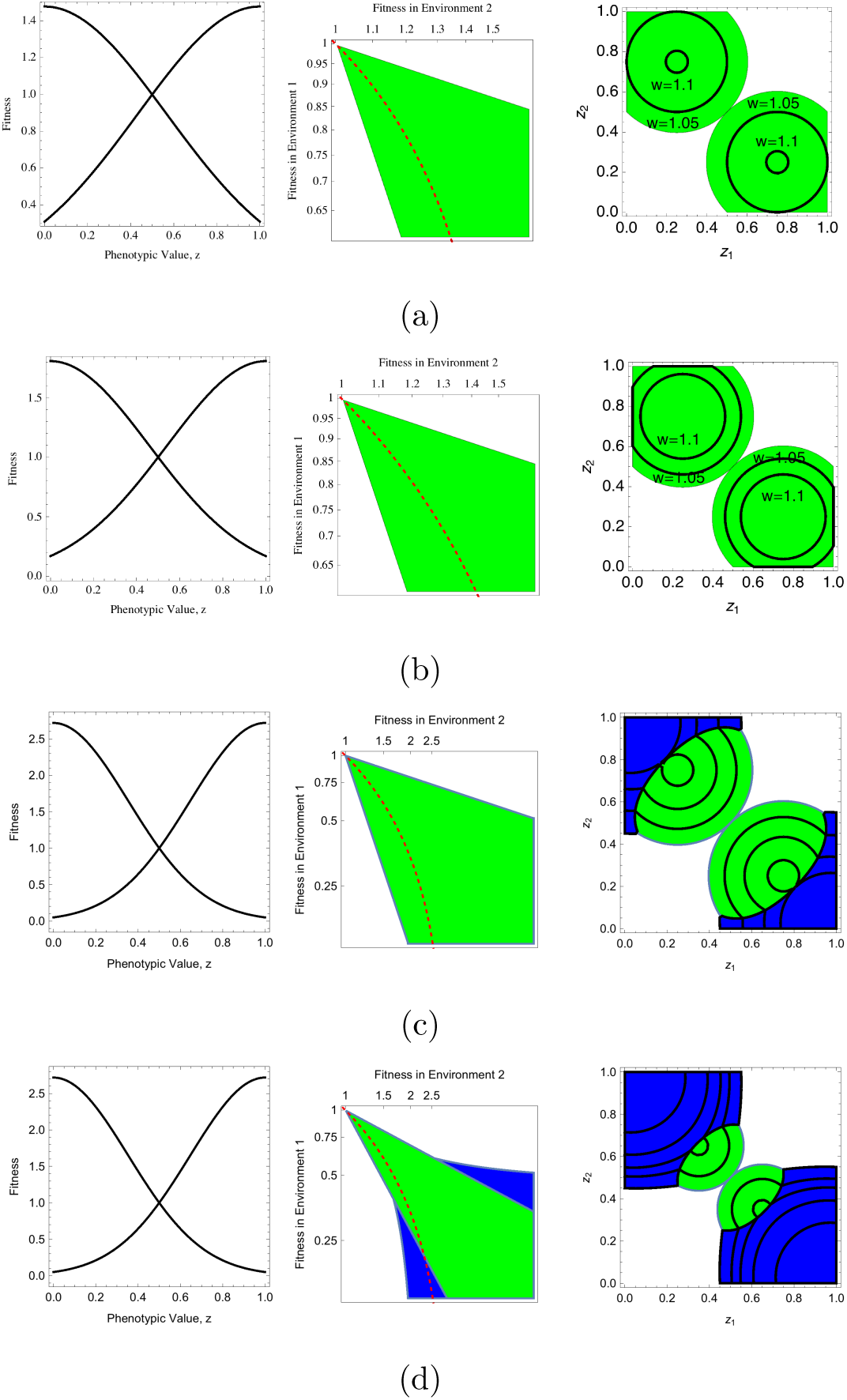
Numerical results for joint evolution of maternal effect strategies and the phenotypes that they control. In panel (a) the parameters are 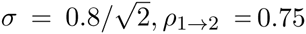, *ρ*_1→2_ = 0.75, *ρ*_2→1_ = 0.75. Under these conditions, RME is never favored over the ancestral, single-phenotype ESS. Panel (b) has parameters 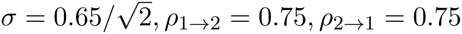, *ρ*_1→2_ = 0.75, *ρ*_2→1_ = 0.75. RME never has higher fitness than DME. Panel (c) has parameters 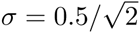, *ρ*_1→2_ = 0.75, *ρ*_2→1_ = 0.75. Panel (d) has parameters 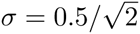, *ρ*_1→2_ = 0.65, *ρ*_2→1_ = 0.65.

When *w* is small enough then RME can invade the single-phenotype (no maternal effect) ESS. In this case, we still expect that DME has higher fitness for mutations that affect *z* by a small amount, but the global behavior may involve evolution of RME. In cases where there is still a large information signal, DME may globally exclude RME. Panel (b) of Figure 10 shows one such example. In this case, the fitness set is concave, but since there is a high degree of predictability to the environment, DME has higher fitness than RME for all phenotype pairs (*z*_1_, *z*_2_).

When *ω* is even lower, RME may have higher values of fitness for some pairs (*z*_1_, *z*_2_) and be the global fitness optimum, and still be unlikely to evolve. Panel (c) of figure 10 shows a case where DME invades near the single-phenotype ESS, and has local maxima at *z*_1_ = 0.75, *z*_2_ = 0.25. RME has highest fitness for the most extreme values of *z*, but has lower fitness in the region of the DME ESS. This suggests that RME may have difficulty evolving even when it is the global optimum.

When both *ω* is lower and the predictability is low, RME may evolve smoothly from DME. Panel (d) of figure 10 shows a case where DME invades near the single-phenotype ESS and RME has higher fitness for *z* values close enough to the single-phenotype ESS that RME can smoothly evolve. The contours show that RME will continue to evolve more extreme values of *z* until the population reaches the global optimum at *z*_1_ = 1, *z*_2_ = 0.

### 3.4 Simulation Results

Our analytical results on the invasion of maternal effect mutants are based on small mutational effects arising near a single phenotype ESS. A simulation approach allows the influence of several processes to be included. Although approximations exist for the probability of fixation in a fluctuating environment (tuljapurkar, 1982; Saether and Engen, 2015), they do not incorporate induced frequency dependence (Proulx, 2000; Lande, 2007, 2008; Proulx and Adler, 2010). Stochastic tunneling may also contribute to the evolution of maternal effects, since multiple substitutions may be required to alter both the phenotypes produced and their frequency of production (Iwasa et al., 2004; Weissman et al., 2009; Lynch and Abegg, 2010; Proulx, 2011). These effects are hard to account for in most analytical formalisms.

We use the Gaussian selection scheme outlined in the previous section with environments having optimum trait values of 0 and 1. Our simulation characterizes individual genotype via parameters that describe the type of maternal effect, the phenotypes produced, and in the case of RME, the probability of producing each phenotype. The first parameter is a discrete variable representing the maternal effect strategy of the individual and can be ‘G’ for a genetically determined phenotype, ‘DME’, or ‘RME’. Mutations alter the type of maternal effect with probability *μ*_ME_ and alter the parameters determining phenotype and probability of producing each phenotype with probability *μ*.

G individuals have only one other parameter, *z*, their phenotype value. Upon reproduction, these individuals can experience mutations in the parameter *z*, following a truncated normal distribution between 0 and 1, with the mutational variance described by 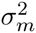. When mutations in the matronal effect strategy itself occur (with probability *μ*_ME_), the maternal effect strategy is switched to one of the other two strategies with equal probability, while the phenotype parameter is maintained at the same value. If the maternal effect strategy changes to RME then the probability of phenotype production is drawn from a uniform distribution.

DME individuals are characterized by two phenotype parameters, *z*_1_ and *z*_2_. For simplicity, *z*_1_ is produced when mothers experience environment 1 and *z*_2_ is produced when mothers experience environment 2. Mutations in either phenotype are equally likely and are again drawn from a normal distribution centered around the current phenotype. If a mutation alters the maternal effect strategy to G, one of the two phenotypes is selected, with equal probability. If a mutation alters the maternal effect strategy to RME, both phenotypes are maintained, and the probability of phenotype production is drawn from a uniform distribution.

RME individuals are characterized by two phenotype parameters, *z*_1_ and *z*_2_ and by *γ*, their probability of producing phenotype *z*_1_. Mutations are equally likely to affect *z*_1_, *z*_2_, and *γ* and are drawn from truncated normal distributions. If a mutation alters the maternal effect type to DME, the two *z* values are maintained. If a mutation alters the maternal effect to G then one phenotype is selected at random as the *z* value.

Figure 11 shows a sample of simulation results for a range of parameters spanning the conditions that are predicted to lead to DME and RME. For fitness functions that are not steep enough, DME readily evolves and the year-to-year variance in phenotype increases. For fitness functions that allow the possibility of RME, if the environmental sequence is predictable enough we still observe DME. When the fitness function is both steep and information content of the environmental sequence is reduced we observe an initial evolution of DME with a narrower range of phenotypes produced. Eventually RME invades and the range of phenotypes increases, but the year-to-year variance in phenotype goes down.

**Figure 11:**
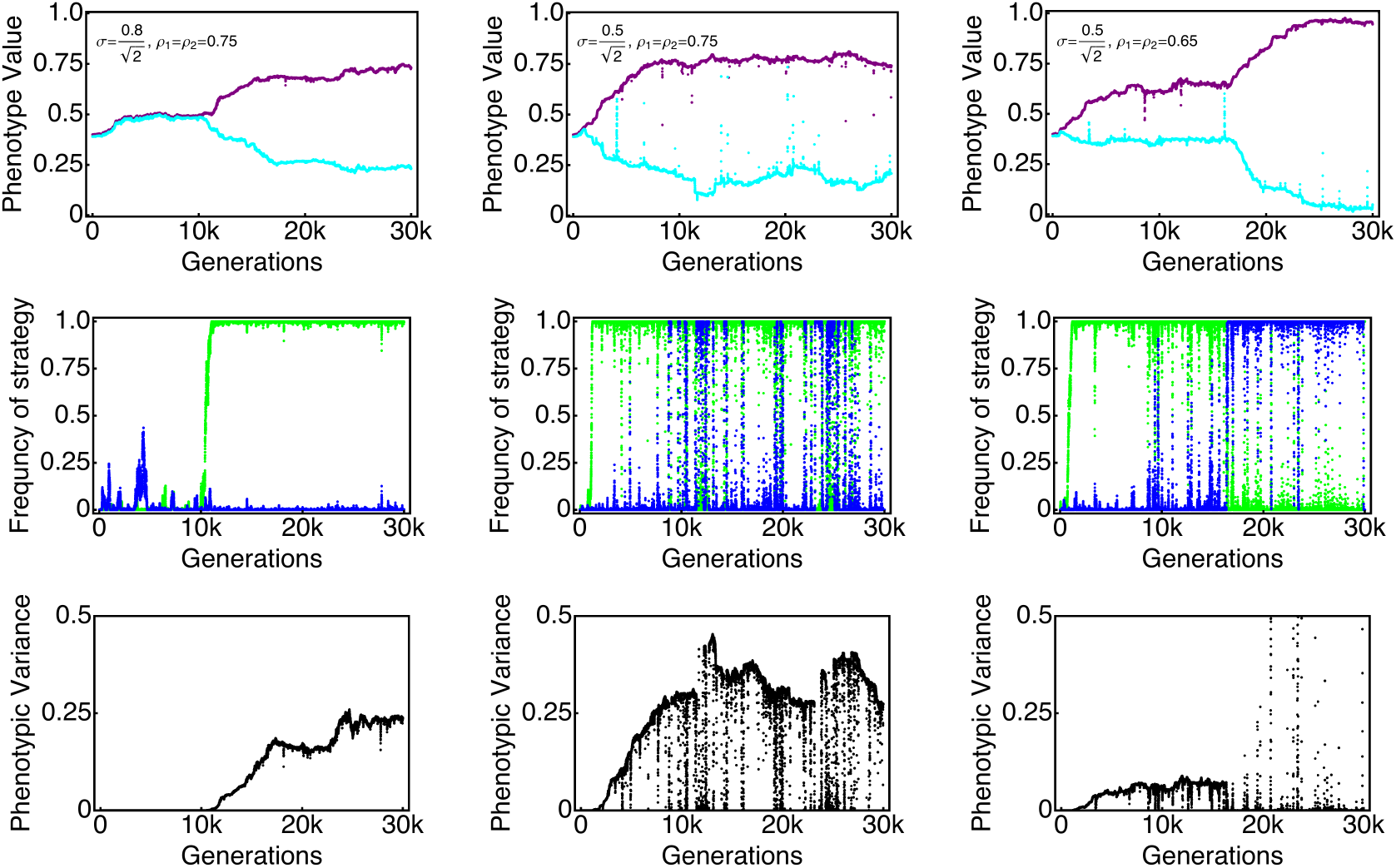
Simulation results: Each column is for different conditions. The top row shows the mean of the two (or one) phenotypes produced (shown in purple and cyan). For individuals that produce a purely genetic strategy, the two phenotypes are constrained to be equal. The middle row is the frequency of aDME (green) and RME (blue). In these simulations mutations altered the phenotype parameter with a probability of 10**^−^**^3^ following a normal distribution with variance of 10^−4^. The maternal effect strategy mutated with probability 10^−4^. The bottom row shows the generation-to-generation variance in phenotype. In Column 1, the parameters are 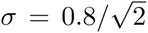, *ρ*_1→2_ = 0.75, *ρ*_2→1_ = 0.75. aDME is expected to evolve first and remain the global ESS. The phenotypic variance increases as aDME evolves and then remains steady. Column 2 has parameters 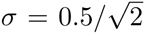, *ρ*_1→2_ = 0.75, *ρ*_2→1_ = 0.75, and aDME is still expected to evolve and be a local ESS. Column 3 has parameters 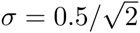, *ρ*_1→2_ = 0.65, *ρ*_2→1_ = 0.65, and RME is expected to invade aDME. Note that aDME first evolves to a quasi-steady state level of phenotypic variance. Once the RME invades, the magnitude of the phenotypic difference increases substantially.

We performed 100 simulations of each set of parameters shown in figure 11 to test these general conclusions. For this batch of simulations, we initialized the populations at the single-phenotype ESS (*z_i_* = 0.5). We then ran the simulation until either DME or RME reached and maintained a frequency of at least 0.95 for 500 generations and the difference between the two phenotypes produced was at least 0.05. The latter requirement was to eliminate situations where a maternal effect mutant that recapitulated the pure genetic strategy became common through drift. For conditions that strongly favor DME (figure 11, column 1), DME evolved in 100 out of 100 times (in less than 40,000 generations). For conditions that have both a DME and RME ESS (figure 11, column 2), DME evolved first in 97/100 times. We continued the simulations in the 3 cases where RME reached high frequency and found that DME replaced RME within 2000 generations. Finally, in situations where we expect DME to evolve first but be replaced by RME (figure 11, column 3), we still found that in 97/100 simulations DME evolved first. In the other 3 simulations RME reached high frequency, and continued simulation resulted in the maintenance of RME in 1 out of 3 cases, but a reversion to DME in the other 2 simulations. Overall, our simulation evidence backs up the idea that DME evolves first and is only replaced by RME when it is strongly favored.

## 4 Conclusions

While there is an extensive theoretical literature on population genetics under variable environmental conditions and selection for maternal effects, our goal here is to use a common theoretical framework to follow the evolution of maternal strategies where the maternal strategy can involve altering offspring phenotype based on the maternal environment. We classify the maternal strategies as deterministic (DME), randomized (RME), or hybrid (HME). Our definitions of maternal effect strategies are based on the mechanism of the maternal effect, and encompass the notions of diversified bet-hedging and anticipatory maternal effects (Philippi and Seger, 1989; Seger and Brockman, 1987; Uller, 2008),without conflating the selective outcome of a maternal effect with the mechanistic basis of that effect. We took two complementary approaches to understanding the evolution of maternal effect, the first looked at the relative advantage of alternative MEs when the set of possible offspring phenotypes was held constant, while the second approach considered the joint evolution of the maternal effect and the phenotypes assuming a continuum of possible phenotypes.

By considering phenotypes with fixed effects we were able to examine the conditions under which the three maternal effect strategies has the largest advantage over a purely genetic strategy. We found that DME is advantageous only when there is predictive power in knowing the maternal environment, while RME is advantageous when the frequency of the two environments falls within a band of intermediate values. The relative fitness effects in the two environments determines the range environmental conditions that favor DME; in contrast increased magnitude of the effects makes RME more advantageous. The environmental parameters that favor RME over DME are in regions where the maternal environments has little predictive information. If producing more offspring with one phenotype entails a fecundity cost, then we find that the range of environmental conditions that favor either DME or RME are shifted, and that relatively small fecundity effects are enough to cause a purely genetic strategy to have highest fitness. While HME always has fitness that is equal or greater than either DME or RME, the HME strategy collapses onto the DME strategy in a wide area of parameter space. Only when the maternal environment provides limited information is an HME strategy the most fit. For environments that have more information available, the HME strategy is to produce a fixed phenotype for each of the possible maternal environments.

We also considered an evolutionary scenario where, in addition to genetic variance in the maternal effect strategy, the set of phenotypes produced could evolve. The idea is that the mutations can introduce a new set of phenotypes produced by the maternal effect. Since the environment-specific fitnesses depend on the phenotype, this also evolves. The results developed in the fixed fitness effect model imply that if the two phenotypes produced are similar to each other then the range of environmental parameters that favor RME will be small. Our general model assumes that before any maternal effect has evolved the population has time to reach the single phenotype ESS. This is a point where a purely genetic strategy is fixed in the population, and all mutants with nearby phenotypes have lower fitness. We then investigated the strength of selection on a maternal effect strategy (either DME or RME) that had access to mutant phenotypes near this ancestral state. We found that mutations that create DME always spread near the ancestral state as long as the maternal environment has some predictive power. In contrast, RME can only invade if the fitness functions are sufficiently steep, an effect that has been previously noted (Bull, 1987). In addition, we found that even when the fitness functions are steep enough to allow RME to invade a purely genetic strategy, mutations that cause DME are always at a selective advantage.

Several authors have explored multiple ways that offspring phenotype can be determined. While we here explored models where the parental genotype and parental environment combine to determine offspring phenotype, other possibilities range from systems where parental phenotype alone determines offspring phenotype to systems where a number of different inputs combine to determine offspring phenotype. The literature on epigenetic effects has focused primarily on cases where parental phenotype determines offspring phenotype (Jablonka et al., 1995), and has been extended to quantitative genetic models of phenotypic inheritance (Hoyle and Ezard, 2012; Kuijper and Hoyle, 2015; Tufto, 2015; Ezard et al., 2014). In these models, an Markovian environmental shift such as we model here can result in the evolution of similar maternal effects as we see here. In particular, these studies found that when the environment tends to alternate between states, a “negative maternal effect” tends to evolve, whereby mothers who survive to breed tend to have phenotypes that will not be adaptive in the next generation, and because of the negative maternal effect these mothers tend to produce offspring with phenotypes that more closely match the expected environment in the next generation (Kuijper and Hoyle, 2015). Such quantitative genetic models, however, explain the maternal effect as a statistical property and do not shed light on the mechanistic basis of the response. Another recent approach has included a range of possible information sources that a developing individual could use as a developmental cue (Leimar and McNamara, 2015), which has the benefit of including both genetic and cue-based phenotypic effects.

Donaldson-Matasci et al. (2013, 2010) considered the fitness value of information and found an upper bound to selection for a generic phenotypically plastic strategy (i.e. any combination of developmental plasticity or parental effect). Their analysis concerns fixed sets of phenotypes, which gives useful insight into the evolutionary dynamics and could, in principal, be combined with a model of phenotype/fitness trade-offs to determine the invasion dynamics of new mutations. Our approach does this without explicitly using information theory to determine the strength of selection on maternal effect mutants, but a combination of the approaches may prove useful.

In this paper we have focused on scenarios with environmental autocorrelation between generations and explored how maternal strategies that alter offspring phenotype evolve. Our modeling framework excludes scenarios where offspring can gain direct information on the environment that they are developing in, and thus limits the scope for the evolution of developmental plasticity. Other work has shown that the relative information content the developing individuals and that parents have can determine whether maternal effects or developmental plasticity evolve (Kuijper and Hoyle, 2015; Leimar and McNamara, 2015), but including opportunities for developmental plasticity to evolve is not likely to make the evolution of RME more likely. If developmental plasticity evolves, either alone or in conjunction with a maternal effect, then the fitness of such a strategy would only increase over a DME strategy, making the invasion of RME even more difficult.

While our results show that the scope for the evolution of RME is low, there are specific conditions that make RME more likely. There must be a large potential fitness benefit of having specialized phenotypes, allowing the evolution of highly specialized phenotypes that have high fitness in a subset of environments and are more or less lethal in other environments.

## 5 Acknowledgements

This work received funding from the National Science Foundation (EF-1137835), the European Research Council (FP7/2007-286 2013/243285), and from the “pepiniere interdisciplinaire” CNRS-PSL Eco-Evo-Devo, the ERC under the European CommunityOs Seventh Framework Program (FP7/2007-2013, Grant Agreement no. 243285).

